# Reduced steroid activation of elephant shark GR and MR after inserting four amino acids from the DNA-binding domain of lamprey corticoid receptor-1

**DOI:** 10.1101/2023.02.24.529974

**Authors:** Yoshinao Katsu, Jiawen Zhang, Michael E. Baker

## Abstract

Atlantic sea lamprey contains two corticoid receptors (CRs), CR1 and CR2, that have identical amino acid sequences, except for a four amino acid insert (Thr-Arg-Gln-Gly) in the CR1 DNA-binding domain (DBD). Steroids are stronger transcriptional activators of CR2 than of CR1 suggesting that the insert reduces the transcriptional response of lamprey CR1 to steroids. The DBD in elephant shark mineralocorticoid receptor (MR) and glucocorticoid receptor (GR), which are descended from a CR, lack these four amino acids, suggesting that a CR2 is their common ancestor. To determine if, similar to lamprey CR1, the presence of this insert in elephant shark MR and GR decreases transcriptional activation by corticosteroids, we inserted these four CR1-specific residues into the DBD of elephant shark MR and GR. Compared to steroid activation of wild-type elephant shark MR and GR, cortisol, corticosterone, aldosterone, 11-deoxycorticosterone and 11-deoxycortisol had lower transcriptional activation of these mutant MR and GR receptors, indicating that the absence of this four-residue segment in the DBD in wild-type elephant shark MR and GR increases transcriptional activation by corticosteroids.

## Introduction

The sea lamprey (*Petromyzon marinus*) belongs to an ancient group of jawless vertebrates known as cyclostomes, which are basal vertebrates that evolved about 550 million years ago (1–4). Sea lamprey contains a corticoid receptor (CR) (5–8), which belongs to the nuclear receptor family of transcription factors (9,10), which also contains the mineralocorticoid receptor, glucocorticoid receptor, progesterone receptor, estrogen receptor and androgen receptor (9–13). A CR in an ancestral cyclostome is the common ancestor to the mineralocorticoid receptor (MR) and the glucocorticoid receptor (GR) in vertebrates (5,7). The MR and GR first appear as separate steroid receptors in cartilaginous fish (5–7,14,15). These two closely related steroid receptors regulate important physiological responses in vertebrates. The MR regulates electrolyte transport in the kidney and colon (12,16–21), as well as regulating gene transcription in a variety of non-epithelial tissues (22–26). The GR has diverse physiological actions in vertebrates including in development, metabolism, the stress response, inflammation and glucose homeostasis (27–31).

The importance of the MR and GR in vertebrate physiology stimulated our interest in understanding the evolution of the MR and GR from the CR (7,8,32). Unfortunately, as a result of complexities in the sequencing and assembly of the lamprey’s highly repetitive and GC rich genome (3,33,34) until recently, only a partial CR sequence was available in GenBank (5).

Fortunately, the recent sequencing of the lamprey germline genome (35) provided contiguous DNA encoding the complete sequences two CR isoforms, CR1 and CR2, which differ only in a novel four amino acid insertion, Thr, Arg, Gln, Gly, (TRQG) in the DNA-binding domain (DBD) (8) (Figure 1).

**Figure 1.**
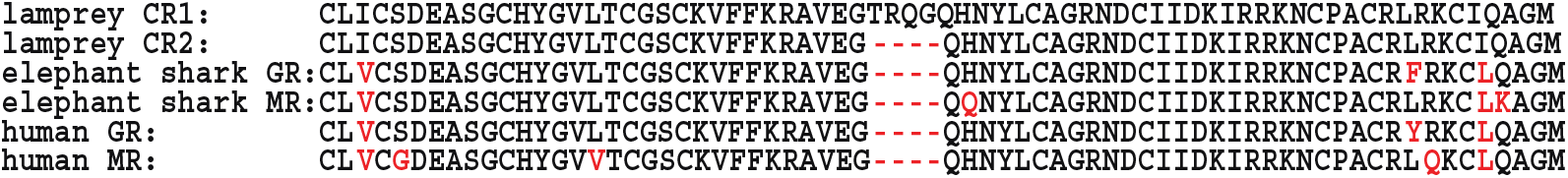
DNA-binding domains of lamprey CR1 and CR2 and elephant shark and human MR and GR. The DNA-binding domain of lamprey CR1 has an insertion of four amino acids that is absent in CR2 and in elephant shark and human MR and GR. Differences between the sequence of lamprey CR and elephant shark and human MR and GR are shown in red.Accessions are: XP_032811370 and XP_032811371 for lamprey CR1 and CR2, respectively, XP_042195980 and XP_007902220 for elephant shark GR and MR, respectively, NP_000167 and AAA59571 for human GR and MR respectively.

With the two CR sequences in hand, we characterized the response of HEK293 cells transfected with each CR isoform to corticosteroids (8) and found that the loss of these four amino acids in the DBD of CR2 increased fold transcriptional activation by about two-fold in the presence of corticosteroids such as 11-deoxycorticosterone, 11-deoxycortisol, cortisol, corticosterone and aldosterone.

Elephant shark mineralocorticoid receptor (MR) and glucocorticoid receptor (GR), which are descended from the CR, lack these four amino acids in their DBD, suggesting that they evolved from a CR2 ancestor. To determine if, similar to lamprey CR2, the absence of this insert in the DBD of elephant shark MR and GR has a functional consequence, we inserted these four residues into their DBDs to convert them to a CR1-like sequence. Here we report that cortisol, corticosterone, aldosterone, 11-deoxycorticosterone and 11-deoxycortisol have lower transcriptional activation of these mutant elephant shark MR and GR receptors, indicating that compared to mutant elephant shark MR and GR, the absence of this four-residue segment in the DBD in both wild-type elephant shark MR and GR increases activation by corticosteroids.

## Results

### Steroid-dependent activation of mutated elephant shark MR and GR

To gain a quantitative measure of steroid activation of mutated elephant shark MR and GR, we determined the concentration dependence of transcriptional activation by corticosteroids of FLAG-tagged wild-type and mutated elephant shark GR transfected into HEK293 cells with either an MMTV-luciferase promoter (Figure 2A-E) (9,36,37) or a TAT3 luciferase promoter (9,38,39) (Figure 2F-J). A parallel study was done with wild-type and mutated elephant shark MR (Figure 3). Luciferase levels were used to calculate an EC50 value and fold-activation for each steroid for FLAG-tagged wild-type and mutated elephant shark GR and MR (Table 1). An EC50 for 11-deoxycortisol activation of FLAG-tagged wild-type GR and elephant shark GR-DBD-TRQG was not determined because fold-activation by 11-deoxycortisol of FLAG-tagged wild-type GR and elephant shark GR-DBD-TRQG did not saturate (Figure 2D, 2J).

**Table 1.**
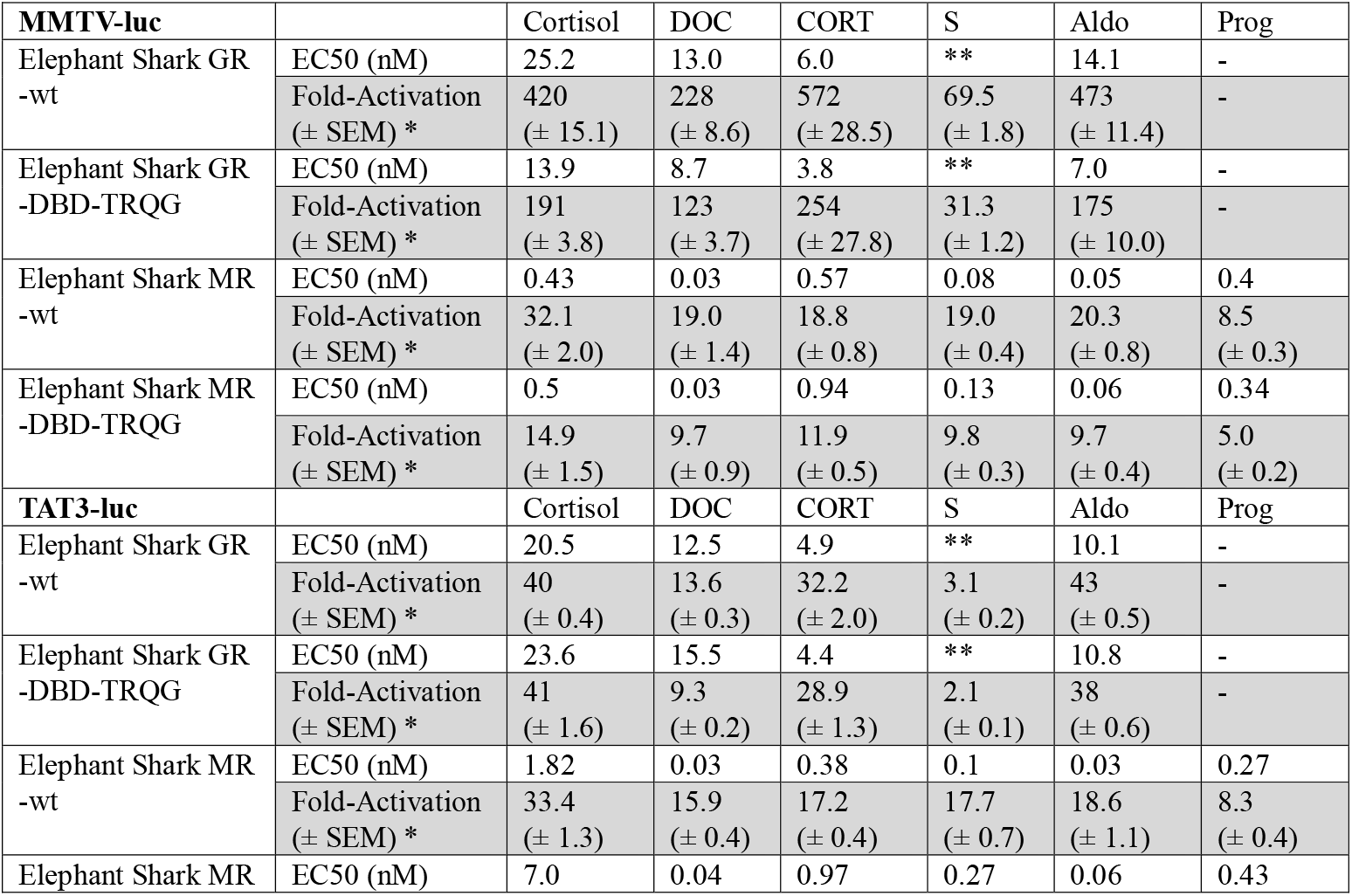
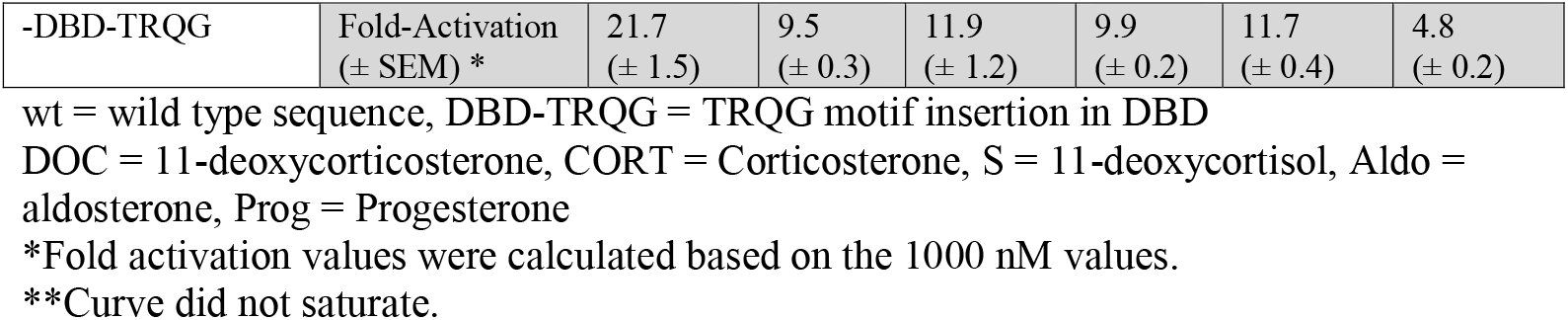
Steroid Activation of FLAG-tagged Wild-Type and Mutated Elephant Shark MR and GR in HEK293 Cells with either an MMTV Promoter or a TAT3 Promoter.

**Fig 2.**
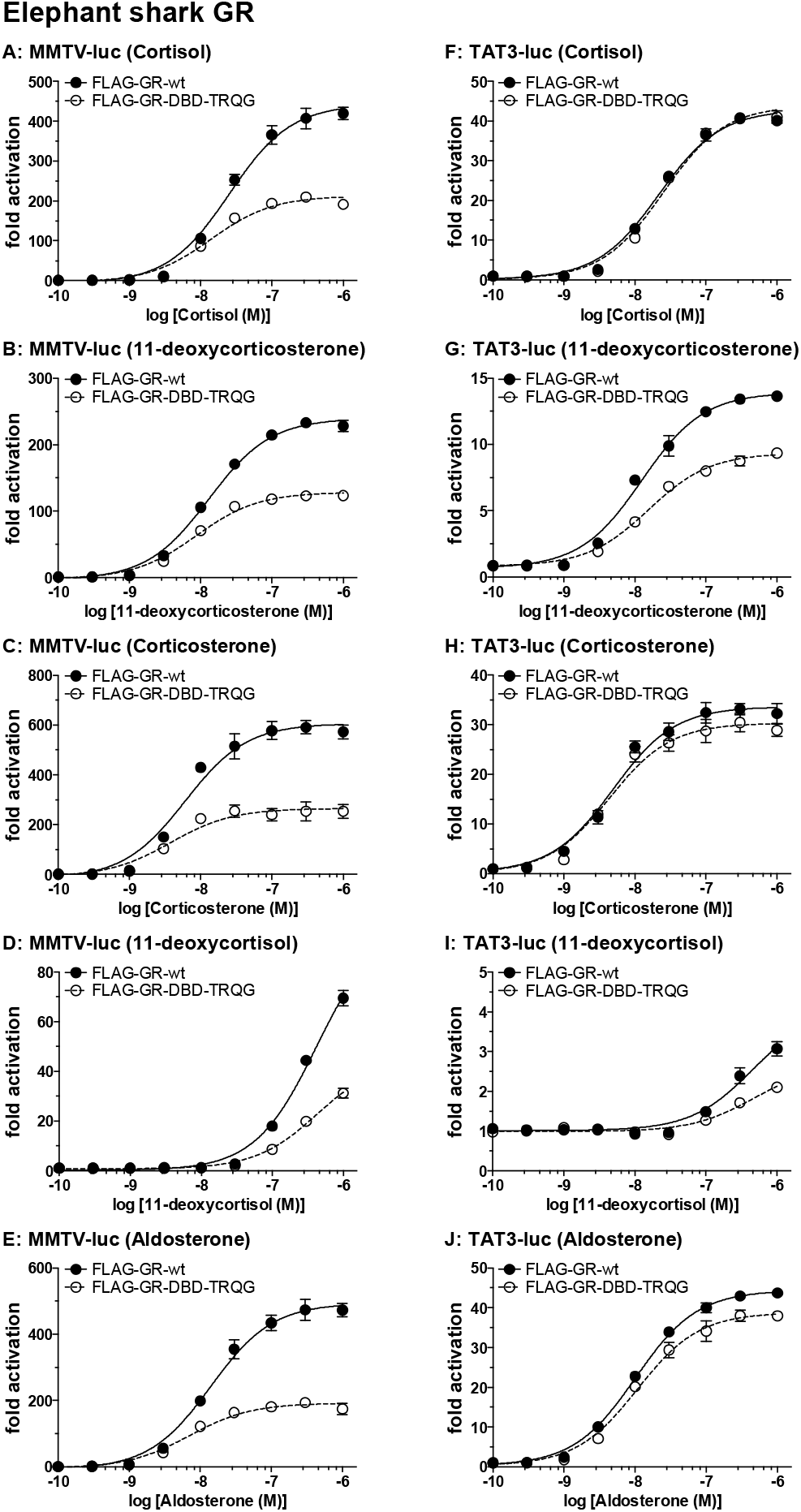
Concentration-dependent transcriptional activation by corticosteroids of FLAG-tagged wild-type and mutated elephant shark GR. Plasmids for FLAG-tagged wild-type elephant shark GR and mutated elephant shark GR (with a TRQG insert) were expressed in HEK293 cells with either an MMTV-luciferase promoter or a TAT3-luciferase promoter (9,36– 38). Cells were treated with increasing concentrations of either aldosterone, cortisol, corticosterone, 11-deoxycortisol, 11-deoxycorticosterone or vehicle alone (DMSO). Results are expressed as means ± SEM, number (n) of wells for each point =3. Y-axis indicates fold-activation compared to the activity of vector with vehicle (DMSO) alone as 1. Figure 2A-E. FLAG-tagged elephant shark GRs with MMTV-luc. Figure 2F-J. FLAG-tagged elephant shark GRs with TAT3-luc.

**Fig 3.**
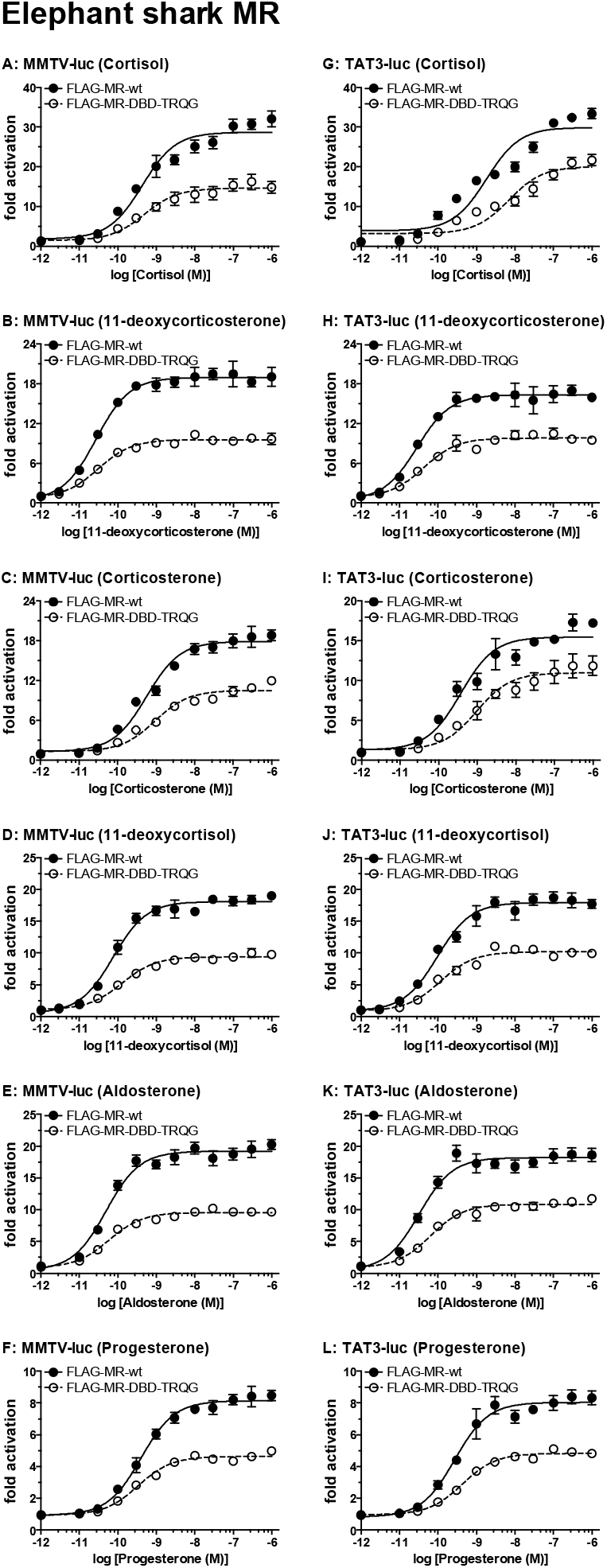
Concentration-dependent transcriptional activation by corticosteroids and progesterone of FLAG-tagged wild-type and mutated elephant shark MR. Plasmids for FLAG-tagged wild-type elephant shark MR and mutated elephant shark MR (with a TRQG insert) were expressed in HEK293 cells with either an MMTV-luciferase promoter or a TAT3-luciferase promoter. Cells were treated with increasing concentrations of either aldosterone, cortisol, corticosterone, 11-deoxycortisol, 11-deoxycorticosterone, progesterone or vehicle alone (DMSO). Results are expressed as means ± SEM, number (n) of wells for each point =3. Y-axis indicates fold-activation compared to the activity of vector with vehicle (DMSO) alone as 1. Figure 3A-F. FLAG-tagged elephant shark MRs with MMTV-luc. Figure 3G-L. FLAG-tagged elephant shark MRs with TAT3-luc.

The results shown in Figure 2 and summarized in Table 1 reveal that addition of the TRQG sequence to the DBD in elephant shark GR reduces fold-activation of this receptor by corticosteroids in HEK293 cells transfected with either the MMTV promoter or the TAT3 promotor. Fold-activation varied depended on the steroid and whether the MMTV promoter or TAT3 promoter was used in the assay. In assays with the MMTV promoter, fold-activation was higher for FLAG-tagged wild-type GR compared to the GR-DBD-TRQG mutant for all tested corticosteroids (Figure 2A-E). Fold-activation also was higher in the assays with the MMTV promoter than with the TAT3 promoter. In assays with the TAT3 promoter, fold-activation was higher for FLAG-tagged wild-type GR compared to the GR-DBD-TRQG mutant for 11-deoxycorticosterone and slightly higher for corticosterone and aldosterone (Figure 2G,H,I). Cortisol had similar fold-activation for FLAG-tagged wild-type GR and GR-DBD-TRQG in assays with TAT3 (Figure 2F). There was little activation by 11-deoxycortisol (Figure 2I).

In assays with the MMTV promoter, cortisol, 11-deoxycorticosterone, corticosterone and aldosterone have a right-shift in the EC50 for FLAG-tagged wild-type elephant shark GR compared to GR-DBD-TRQG (Table 1, Figure 2A-2C, 2E). In the presence of the TAT3 promoter, 11-deoxycorticosterone has a small right shift in the EC50 for FLAG-tagged wild-type GR compared to GR-DBD-TRQG (Table 1, Figure 2G). In the presence of the TAT3 promoter, cortisol (Figure 2F), corticosterone (Figure 2G) and aldosterone (Figure 2J) have similar EC50s for FLAG-tagged wild-type elephant shark GR compared to GR-DBD-TRQG. Corticosteroids also had lower transcriptional activation of FLAG-tagged elephant shark MR-DBD-TRQG compared to FLAG-tagged wild-type elephant shark MR (Figure 3, Table 1).

FLAG-tagged elephant shark MR and elephant shark MR-DBD-TRQG also had higher fold-activation with MMTV promoter compared to the TAT3 promoter, although it varied with the corticosteroid. For example, fold-activation of DOC for elephant shark MR was 12.4 in the presence of MMTV and 8.0 in the presence of TAT3. And fold-activation of aldosterone for elephant shark MR was 12 in the presence of MMTV and 10.7 in the presence of TAT3.

We also studied progesterone activation of FLAG-tagged elephant shark MR with TRQG in its DBD. Progesterone is an antagonist for human MR (40–42), and an agonist for fish MR (14,41,43,44), Progesterone is a transcriptional activator of elephant shark MR, but not elephant shark GR (14,45). We find that addition of the TRQG sequence to FLAG-tagged elephant shark MR reduces fold-activation by progesterone in HEK293 cells transfected with either the MMTV promoter or the TAT3 promotor (Figure 3).

### FLAG-tagged GR and MR protein expression

To confirm the protein expression levels of FLAG-tagged GR and MR cultured cells transfected with each construct were collected and treated with SDS sample buffer. And then, 20 μg of protein was subjected to SDS-PAGE on a 10% polyacrylamide gel and transferred to an Immobilon membrane. The gels were also Coomassie blue, CBB stained. Subsequently, FLAG-tagged proteins were detected using an antibody against the FLAG tag. We detected a single band the MR and multiple bands the GR (Figure 4). Both MR and GR showed no marked difference in protein levels between the wild type and the mutant containing 4 amino acid insertion in the DBD. The multiple bands in GR may be due to protein phosphorylation.

**Figure 4.**
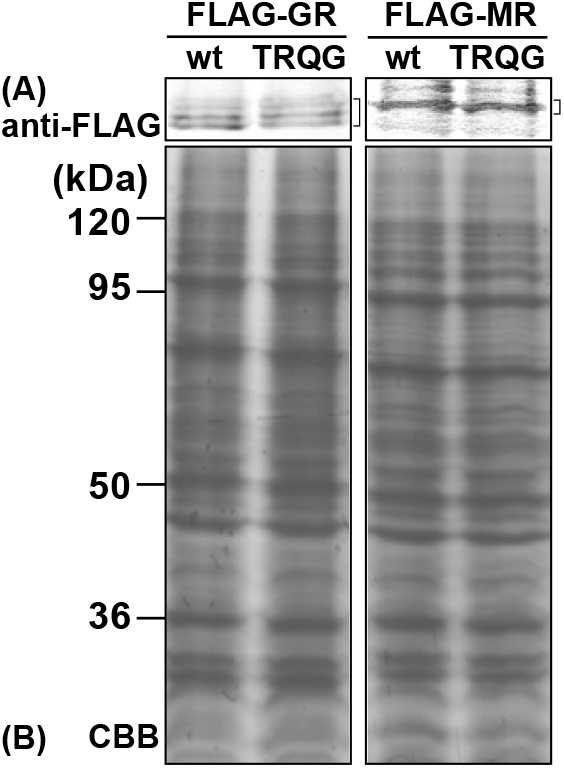
Expression of FLAG-tagged elephant shark GR and MR. HEK293 cell lysates transfected FLAG-tagged constructs were treated with sample buffer and applied to a 10% SDS-polyacrylamide gel, and then transferred onto a membrane. The expressed FLAG-tagged proteins were detected with anti-FLAG antibody. (A) FLAG-tagged elephant shark GR or MR. (B) CBB stain.

## Discussion

The sequencing of lamprey germline genome (35) led to the discovery of two CRs, which differ only in a small four residue deletion in the DBD of CR2 (8). In lamprey this deletion in CR2 increases transcriptional activation by corticosteroids. Both elephant shark MR and GR lack this four amino acid insert in their DBDs, which raised the question of whether adding this four-residue insert to elephant shark MR and GR would reduce corticosteroid-mediated transcription, as seen in lamprey CR1 (8) because, in contrast to lamprey CR1 and CR2, elephant shark MR and GR have different sequences in the LBD (61% identical to each other) and NTD (21% identical to each other). Both domains are important in their response to corticosteroids (14). Thus, these sequence differences could obscure the effect of the four-residue insert in the DBD of elephant shark MR and GR. However, as reported here (Figures 2 and 3, Table 1), we find that like lamprey CR1, addition of four residues -TRQG-to wild-type elephant shark MR and GR leads to a reduction in transcriptional activation by a panel of corticosteroids consisting of cortisol, corticosterone, aldosterone, 11-deoxycorticosterone, 11-deoxycortisol and progesterone. This effect of the TRQG insert in elephant shark MR and GR is found for HEK293 cells transfected with either the MMTV or TAT3 promoters (Table 1). The loss of the TRQG insert in elephant shark GR and MR appears to have provided increased transcriptional activity for their GR and MR.

Although elephant shark MR lacks an insert in the DBD, a four amino acid (KCSW) insert is present at this site in the DBD in human MR (46–48), rat MR (46) and *Xenopus laevis* MR (Figure 5). The insertion of KCSR in the DBD of *Xenopus laevis* MR (Figure 4) indicates that introduction of a mutation at this position occurred in a basal terrestrial vertebrate.

**Figure 5.**
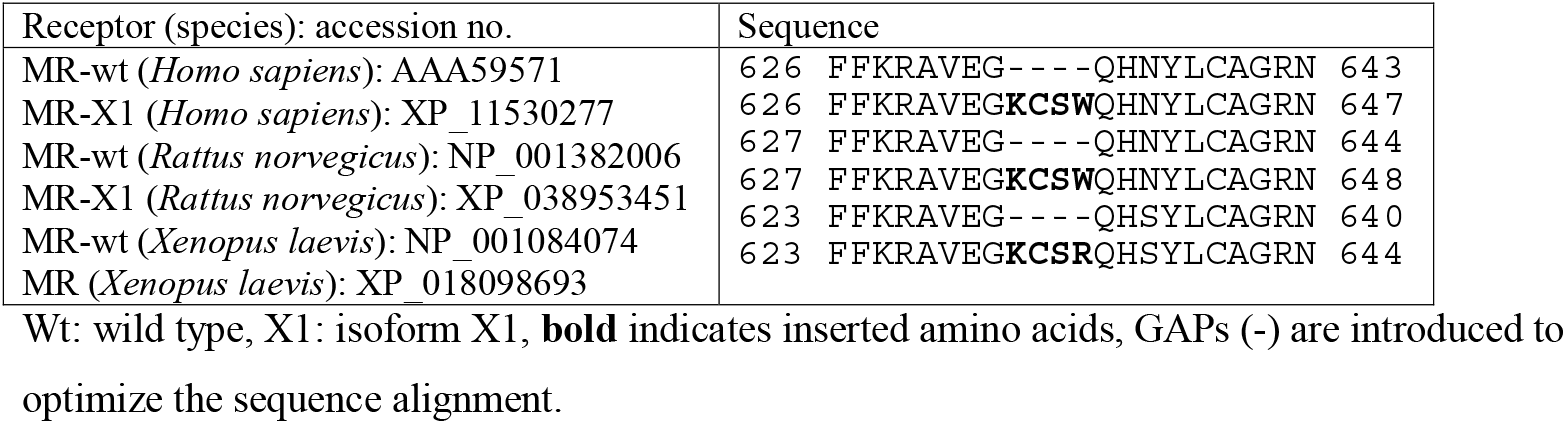
Alignment of Isoforms of the Human, Rat and Frog Mineralocorticoid Receptor DNA-Binding Domain.

Transcriptional activation of these terrestrial vertebrate MRs by steroids has not been reported. We are beginning to investigate corticosteroid and progesterone activation of the human MR variant.

Meijsing et al (49) found that the function of the DBD in human GR was more than just docking of the GR to DNA. They showed that the human GR DBD also had an allosteric effect on transcription by human GR. Our finding that the DBD in elephant shark MR and GR and in sea lamprey CR has a role in regulating corticosteroid-mediated transcription indicates that a regulatory function of the DBD appeared early in the evolution of corticosteroid receptors.

## Methods

### Construction of plasmid vectors

Full-length glucocorticoid receptor (GR) and mineralocorticoid receptor (MR) sequences of elephant shark, *Callorhinchus milii*, were registered in Genbank (accession number: XP_042195980 for GR and XP_007902220 for MR). The insertion of 4-amino acids (Thr-Arg-Gln-Gly) into the DBD of elephant shark MR and GR was performed using KOD-Plus-mutagenesis kit (TOYOBO). N-terminal FLAG-tag motif was introduced into both wild type and mutants containing the 4-amino acid insert in the DBD using PCR. The nucleic acid sequences of all constructs were verified by sequencing.

### Western blotting

FLAG-tagged GR and MR constructs were transfected as described in “Transactivation assays and statistical analysis” (50). FLAG-tagged proteins were separated by SDS-PAGE in 10 % gel, blotted onto an Immobilon membrane (Millipore Corp. MA), and probed with anti-FLAG (clone 2H8, TransGenic Inc. (50)) antibody.

### Chemical reagents

Cortisol, corticosterone, 11-deoxycorticosterone, 11-deoxycortisol, aldosterone and progesterone were purchased from Sigma-Aldrich. For reporter gene assays, all hormones were dissolved in dimethyl-sulfoxide (DMSO); the final DMSO concentration in the culture medium did not exceed 0.1%.

### Transactivation assays and statistical analyses

Transfection and reporter assays were carried out in HEK293 cells, as described previously (8). The cells were transfected with 100 ng of receptor gene, reporter gene containing the *Photinus pyralis* luciferase gene and pRL-tk, as an internal control to normalize for variation in transfection efficiency; pRL-tk contains the *Renilla reniformis* luciferase gene with the herpes simplex virus thymidine kinase promoter. Each assay had a similar number of cells, and assays were done with the same batch of cells in each experiment. All experiments were performed in triplicate. Promoter activity was calculated as firefly (*P. pyralis*)-lucifease activity/sea pansy (*R. reniformis*)-lucifease activity. The values shown are mean ± SEM from three separate experiments, and dose-response data, which were used to calculate the half maximal response (EC50) for each steroid, were analyzed using GraphPad Prism. We note that our assays were done at 37 degrees, which is not the physiological temperature for the GR and MR in elephant sharks.

## Author Contributions

**Conceptualization:** Yoshinao Katsu, Michael E. Baker.

**Data curation:** Yoshinao Katsu, Jiawen Zhang

**Formal analysis:** Yoshinao Katsu, Michael E. Baker.

**Investigation:** Yoshinao Katsu, Jiawen Zhang

**Methodology:** Yoshinao Katsu.

**Supervision:** Yoshinao Katsu, Michael E. Baker.

**Writing – original draft:** Yoshinao Katsu, Michael E. Baker.

**Writing – review & editing:** Yoshinao Katsu, Michael E. Baker.

